# The secreted acid phosphatase domain-containing GRA44 from *Toxoplasma gondii* is required for C-myc induction in infected cells

**DOI:** 10.1101/867614

**Authors:** William J Blakely, Michael J Holmes, Gustavo Arrizabalaga

## Abstract

During host cell invasion, the eukaryotic pathogen *Toxoplasma gondii* forms a parsitophorous vacuole to safely reside within, while partitioned from host cell defense mechanisms. From within this safe niche parasites sabotage multiple host cell systems including gene expression, apoptosis and intracellular immune recognition by secreting a large arsenal of effector proteins. Many parasite proteins studied for active host cell manipulative interactions have been kinases. Translocation of effectors from the parasitophorous vacuole into the host cell is mediated by a putative translocon complex, which includes proteins MYR1, MYR2, and MYR3. Whether other proteins are involved in the structure or regulation of this putative translocon is not known. We have discovered that the secreted protein GRA44, which contains a putative acid phosphatase domain, interacts with members of this complex and is required for host cell effects downstream of effector secretion. We have determined GRA44 is processed in a region with homology to sequences targeted by protozoan proteases of the secretory pathway and that both major cleavage fragments are secreted into the parasitophorous vacuole. Immunoprecipitation experiments showed that GRA44 interacts with a large number of secreted proteins included MYR1. Importantly, conditional knockdown of GRA44 resulted in a lack of host cell cMyc upregulation, which mimics the phenotype seen when members of the translocon complex are genetically disrupted. Thus, the putative acid phosphatase GRA44 is crucial for host cell alterations during *Toxoplasma* infection and is associated with the translocon complex which *Toxoplasma* relies upon for success as an intracellular pathogen.

**IMPORTANCE:** Approximately one third of humans are infected with the parasite *Toxoplasma gondii*. *Toxoplasma* infections can lead to severe disease in those with a compromised or suppressed immune system. Additionally, infections during pregnancy present a significant health risk to the developing fetus. Drugs that target this parasite are limited, have significant side effects, and do not target all disease stages. Thus, a thorough understanding of how the parasite propagates within a host is critical in the discovery of novel therapeutic targets. To replicate *Toxoplasma* requires to enter the cells of the infected organism. In order to survive the environment inside a cell, *Toxoplasma* secretes a large repertoire of proteins, which hijack a number of important cellular functions. How these *Toxoplasma* proteins move from the parasite into the host cell is not well understood. Our work shows that the putative phosphatase GRA44 is part of a protein complex responsible for this process.

## INTRODUCTION

*Toxoplasma gondii* is an obligate intracellular eukaryotic pathogen infecting an estimated one-third of the human population globally. Approximately 15% of the US population is positive for *Toxoplasma* infection (1), while some countries in Europe and South America have much higher infection rates. Within the human host, *Toxoplasma* exists as either highly-proliferative tachyzoites, which are responsible for the acute stage of the infection, or latent bradyzoite cysts, which form in various tissues and establish a chronic infection. While most infections are asymptomatic, in immunocompromised individuals and lymphoma patients, new infections or re-activation of pre-existing cysts can lead to toxoplasmic encephalitis, among other complications (2–4). Additionally, a primary infection puts pregnant women at risk of passing parasites to the developing fetus, which can cause miscarriage and severe birth defects (5).

Due to its biological niche as an obligate intracellular parasite, *Toxoplasma* depends upon remodeling the host cell environment to facilitate its own growth and survival. As the parasite invades a host cell, it forms an insular vacuole, known as the parasitophorous vacuole (PV), within which they safely replicate undisturbed by host cell innate immune machinery. Within the PV, parasites interface with the host cell through the PV membrane while avoiding direct contact with host cell components. Mediation of interactions between parasites and the host cell is accomplished by a multitude of parasite proteins that are secreted during invasion from the rhoptries (ROP proteins) and during intracellular growth from the dense granules (GRA proteins). Many of these proteins are secreted beyond the PV into the host cell to directly manipulate host processes like transcription, apoptosis, immune responses and metabolism (6, 7). Additional proteins are secreted but retained within the PV for the purpose of trafficking these effectors to the host cytoplasm. The proteins MYR1, MYR2 and MYR3, components of a putative translocon system, are secreted to the PV and are responsible for altering a wide array of host processes by trafficking effectors to the host (8). Secreted proteins, especially those with enzymatic activity such as kinases and phosphatases, are of great interest, as they hold potential as drug targets due to their exclusivity to apicomplexans and importance to parasite survival. Two kinases secreted during invasion and intracellular growth that are known to be critical for host manipulation and parasite virulence are ROP16 (9) and ROP18 (10). The ROP16 kinase acts to downregulate STAT3/6 in the host nucleus, which results in altered transcription (11). ROP18, in partnership with GRA7, counteracts host cell immune responses by phosphorylation and inactivation of host Immunity-Related GTPase (IRG) proteins, which otherwise act to signal PV degradation (12). Other *Toxoplasma* kinases known to be secreted into the host cell include WNG1 and WNG2, formerly ROP35 and ROP34 respectively (13). Additionally, several pseudokinases have been shown to be secreted into the host and are implicated in host-cell manipulation such as ROP5, which complexes with ROP18 conferring binding affinity to host IRGA6 (12).

Whether secreted phosphatases play a similarly important role as do secreted kinases and pseudokinases remains largely unexplored. The secreted phosphatase PP2C has been proposed to reduce apoptosis of infected host cells (14) and the phosphatase PP2C-hn has been found in the host nucleus (15), although its function remains unclear. To expand our understanding of phosphatases secreted to either the host or PV, we bioinformatically identified 32 proteins predicted to have both a phosphatase domain and signal sequence. Of those identified, we characterized the biological role of TGGT1_228170 which was previously identified as part of the inner membrane complex and named IMC2A by Mann and Beckers (16). However, contrary to the initially documented localization, this protein contains characteristics of secreted proteins namely a signal sequence and predicted TEXEL motifs, protease cleavage sites found in many *Toxoplasma* secreted proteins (17). Here we show that this protein is both processed and secreted into the PV where it interacts with the proposed translocation complex of MYR1/2/3. Importantly, we show that TGGT1_228170, now renamed GRA44, is critical for activation of the host oncogenic factor cMYC.

## RESULTS

### Bioinformatic search for secreted phosphatase

To begin identifying putative secreted phosphatases, we used a bioinformatics approach starting with all proteins annotated in the *Toxoplasma gondii* genome database ToxoDB (toxodb.org). A blast search of all *Toxoplasma* genes, filtered by only including genes whose products contain predicted phosphatase domains and signal peptides, generated a list containing 32 proteins of potential interest (Table S1 in supplemental material). To prioritize our studies on phosphatases that are likely to play an important role in parasite propagation, we ranked our list according to gene fitness scores as assigned through a genome-wide CRISPR/Cas9 knockout study (18). Among these is TGGT1_228170, a protein that contains a predicted acid phosphatase domain (Fig. 1A) and, that despite having a signal sequence, was previously described as localizing to the inner membrane complex (IMC) (16). However, multiple lines of evidence suggest that it may indeed be secreted. First, the homologous protein UIS2 in the related apicomplexan parasite *Plasmodium berghei* has been shown to have a secreted ortholog (Pf3D7_1464600) in *Plasmodium falciparum* (19). Second, TGGT1_228170 has been repeatedly detected in BioID experiments as an interactor of proteins localized to the PV lumen and PV membrane microenvironments (20). Finally, analysis of the protein sequence revealed multiple putative ***Toxoplasma* Ex**port **El**ements (TEXEL), which at the time this investigation began was defined as RxLxD/E (21), and has since been refined as RRL (22). TEXEL sequences are recognized by a Golgi-associated protease, aspartyl protease V (ASP5), which cleaves proteins as part of the secretory pathway to the PV/PVM and host cell. Based on these criteria and the fact TGGT1_228170 was assigned a gene knockout fitness score of −3.28, indicating that it substantially contributes to parasite fitness, we decided to revisit the localization and function of this protein.

**Figure 1.**
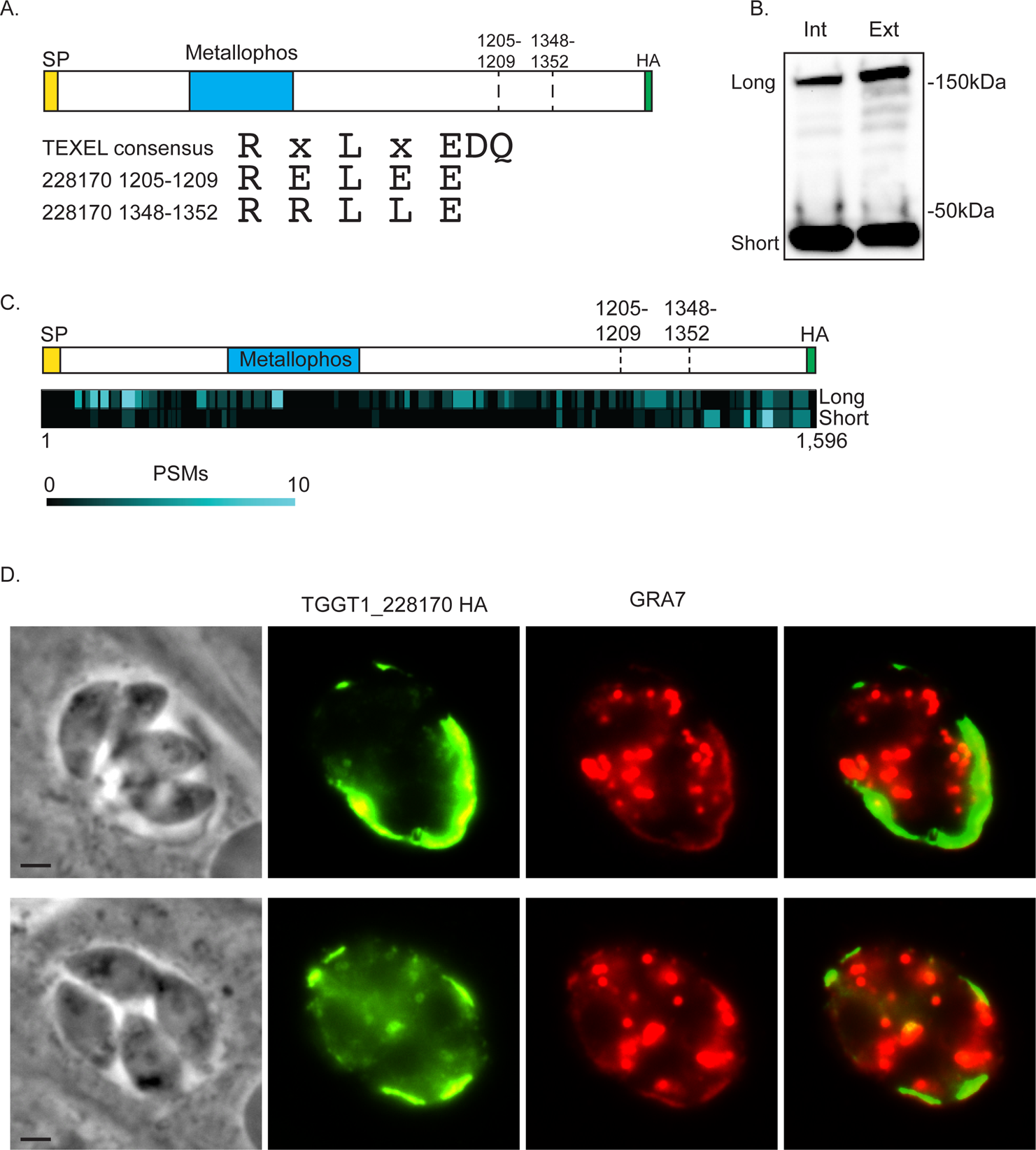
TGGT1_228170 is processed and secreted into the parasitophorous vacuole. To determine the localization of TGGT1_228170 we introduced a C-terminal HA epitope tag in the endogenous gene. A) Schematic showing relative position of the signal peptide (SP), the phosphatase domain (Metallophos), and the two putative TEXEL cleavage sites, 1205-1209 and 1348-1352. The C terminal HA epitope tag is also shown. Below the schematic are the sequences of the TEXEL1 and TEXEL2 cleavage domains compared to the consensus of the *Plasmodium* PEXEL domain. B) Western blot of protein lysates from intracellular and extracellular parasite of the TGGT1_228170(HA) expressing strain probed with antibodies against HA. Two stable forms, labeled as long and short, are detected. C) Heat map illustrating relative position of peptide to spectrum matches (PSMs) from mass spectrometric analysis of long and short forms of TGGT1_228170(HA) in respect to the full protein and the putative cleavage sites. D) Immunofluorescence assay (IFA) of intracellular parasites of the IFA images of strain expressing TGGT1_228170(HA) stained for the parasitophorous vacuole protein Gra7 (in red) and for HA (in green). Scale bar = 2 µm.

### TGGT1_228170 is processed and secreted into the parasitophorous vacuole

To determine the localization of TGGT1_228170, we introduced sequences encoding three C-terminal hemagglutinin epitope tags (3xHA) into the endogenous gene by homologous recombination (Fig. 1A). Western blot analysis of protein extract from both intracellular and extracellular parasites of the TGGT1_228170(HA) line showed a band of approximately 180kDa, which is the expected size for the full protein (Fig. 1B). However, a second prominent band at approximately 40kDa was also noted in both intracellular and extracellular parasites (Fig. 1B). This second smaller band is consistent with processing at either of two areas with homology to TEXEL sites (Fig. 1A). The sequence for the first of these putative cleavage sites is RELEE (amino acids 1205-1209), which is consistent with previous TEXEL consensus, while the sequence for the second is RRLLE (amino acids 1348-1352), which is consistent with the RRL consensus sequencee (Fig. 1A).

To confirm the identity of the 40kDa fragment observed in western blot, endogenously tagged TGGT1_228170 was immunoprecipitated and the eluate separated by SDS-PAGE. Both the 180kDa (long) and 40kDa (short) bands were excised from the PAGE gel and analyzed separately by mass spectrometry (M/S). Results confirmed both bands corresponded to TGGT1_228170. For the long form, we detected 192 peptides distributed throughout the full protein sequence (Fig. 1C). M/S analysis of the band migrating to 40kDa revealed 74 peptides of which 60 were located after the second putative cleavage site(Fig. 1C). Thus, TGGT1_228170 is processed and both the full-length and C-terminal forms are stable.

Finally, to determine the localization of TGGT1_228170, we performed immunofluorescence assays (IFA) of TGGT1_228170(HA) parasites. Consistent with the presence of a signal sequence and putative TEXEL sites, TGGT1_228170 was detected within the PV lumen and at the PV membrane (PVM; Fig. 1D). Based on this localization and corroborative findings by Coffey et al. on the same protein (13), TGGT1_228170 should be designated as a GRA protein, and we will henceforth refer to it as GRA44.

### Amino acids 1348-1352 are required for efficient processing of GRA44

We hypothesized that the small C-terminal fragment of GRA44 detected by Western blot and M/S is the product of cleavage at either of the two putative TEXEL sequences. To investigate which of the two sites is actively cleaved, we exogenously expressed GRA44 with the first arginine of either or both of these sites mutated to alanine (Fig. 2A). The first arginine of TEXEL sites has been previously shown to be important for cleavage (21). Western blot analysis showed that mutating the first arginine of the first putative site (GRA44 R1205A) did not affect processing of the protein (Fig. 2B). By contrast, mutating the arginine in the second site (GRA44 R1348A) significantly reduced processing (Fig. 2B). The same result was observed when both putative TEXEL sites were mutated (GRA44 R1205A/R1348A) (Fig. 2B). Densitometry analysis showed that while for the wild type protein 82.9%±9.9 (n=3) of total protein is cleaved, cleavage in the GRA44 R1348A is 47.4%±9.9 (n=3). Thus, it appears that the second TEXEL is the site for processing of GRA44, and it is referred to as the GRA44 TEXEL from now on. For a thorough examination of the identified TEXEL, we generated parasites exogenously expressing GRA44 in which either L1350 or E1352 are mutated for alanine (Fig. S1A in supplemental material). As was the case for the R1348A mutation, changing the central leucine in the GRA44 TEXEL to an alanine disrupted processing (Fig. S1B in supplemental material), however mutant E1352A showed similar levels of processing as the wildtype protein (Fig. S1B in supplemental material).

**Figure 2.**
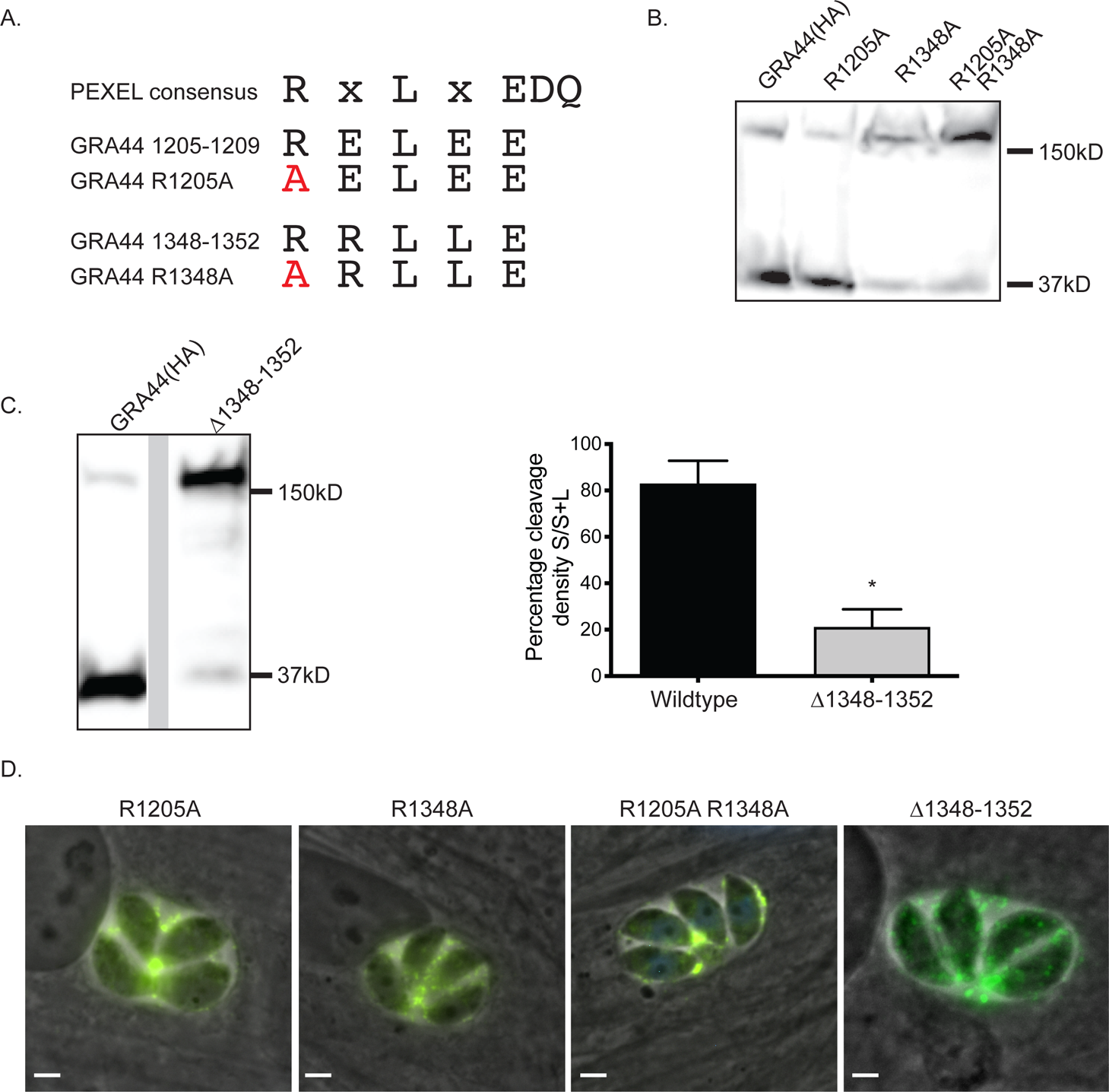
The second putative TEXEL motif is critical for efficient processing of GRA44. To determine the site responsible for the processing of GRA44 we expressed exogenous copies of GRA44(HA) in which the first arginine of the putative cleavage sites were mutated to alanine individually (R1205A and R1348A) or in combination (R1205A/R1348A) or in which the five amino acids of the second putative site were deleted (Δ1348-1352). A) Diagram of the PEXEL consensus and the putative sites in GRA44 with the mutated amino acids indicated in red. B) Western blot of lysates from parasites expressing the exogenous GRA44(HA) or mutant versions R1205A, R1348A and the double mutant R1205A/R1348A probed for HA. C) Western blot of lysates from parasites expressing an exogenous copy of GRA44 lacking the second site (Δ1348-1352) probed with antibodies against the HA epitope tag. Graph represents the percent cleavage in each strain, which was determined by calculating the ratio of the density of the large band over the sum of the density of both bands. n=3, ±SD, *p<0.05 paired t test. D) Representative IFA images of intracellular parasites expressing each of the four GRA44 mutant TEXELs. Scale bars = 2 µm.

Although we observed a significant reduction in cleavage after altering single amino acids in the GRA44 TEXEL, there appeared to be some residual C-terminal cleavage product present with all mutants (Fig. 2B and S1B). To ascertain whether this was the effect of persistent cleavage at TEXEL despite the mutations or cleavage at an alternative site, we generated an exogenously expressed GRA44 mutant in which all five amino acids that make up this TEXEL were deleted (Δ1348-1352). As expected, deletion of the TEXEL resulted in significant loss of the C-terminal cleavage product (Fig. 2C). However, it was evident there still remained a measurable amount being cleaved. Cleavage level for Δ1348-1352 was calculated at 21.1±7.7 (n=3), which is significantly lower than the point mutant R1348A (see above). Since the TEXEL was not present in this mutant form of GRA44, it is plausible that a cryptic site is used, potentially amino acids 1205-1209. Alternatively, GRA44 may be cleaved by a TEXEL/ASP5 independent mechanism.

To determine whether effective processing is needed for localization of GRA44 to the PV, we performed IFAs with parasites expressing each of the four mutant GRA44 (R1205A, R1348A, R1205A/R1348A, and Δ1348-1352). Interestingly, none of the mutations affected secretion and localization to the parasitophorous vacuole (Fig. 2D). Similarly, mutating L1350 or E1352 within the confirmed TEXEL site did not affect the PV localization of GRA44 (Fig. S1C in supplemental material). These results indicate that complete processing is not required for protein secretion.

### Both the N-terminal and C-terminal GRA44 cleavage products localize within the PV

As the localization analysis performed depended on a C-terminal HA epitope tag and GRA44 is cleaved at an internal TEXEL site, we could only detect the full-length uncleaved protein and smaller C-terminal fragment. Consequently, with the C-terminal HA tagged protein we cannot determine the localization of N-terminal cleavage product, which contains the putative acid phosphatase domain. Accordingly, we engineered a strain exogenously expressing GRA44 containing a MYC epitope tag inserted between amino acids 1203 and 1204 in addition to the HA epitope tag at the C-terminus (Fig. 3A). Protein extracts from parasites expressing the dually tagged GRA44 were analyzed by western blot, and probed separately with antibodies against the MYC or HA epitopes (Fig. 3B). Probing anti-MYC uncovered a band at approximately 140kDa in addition to the 180kDa full length protein (Fig. 3B). This 140 kDa correlates to the expected N-terminal end of GRA44 post-cleavage. As observed previously, probing with antibodies against the HA epitope revealed the full-length GRA44 and the C-terminal fragment. Having established a parasite line that allows monitoring of two post-processing fragments of GRA44, we investigated their respective localization by IFA. Regardless of whether we used HA or MYC antibodies, the protein was localized primarily to the PV, which suggests that the two major fragments, including the one containing the phosphatase domain, are secreted into the parasitophorous vacuole (Fig. 3C).

**Figure 3.**
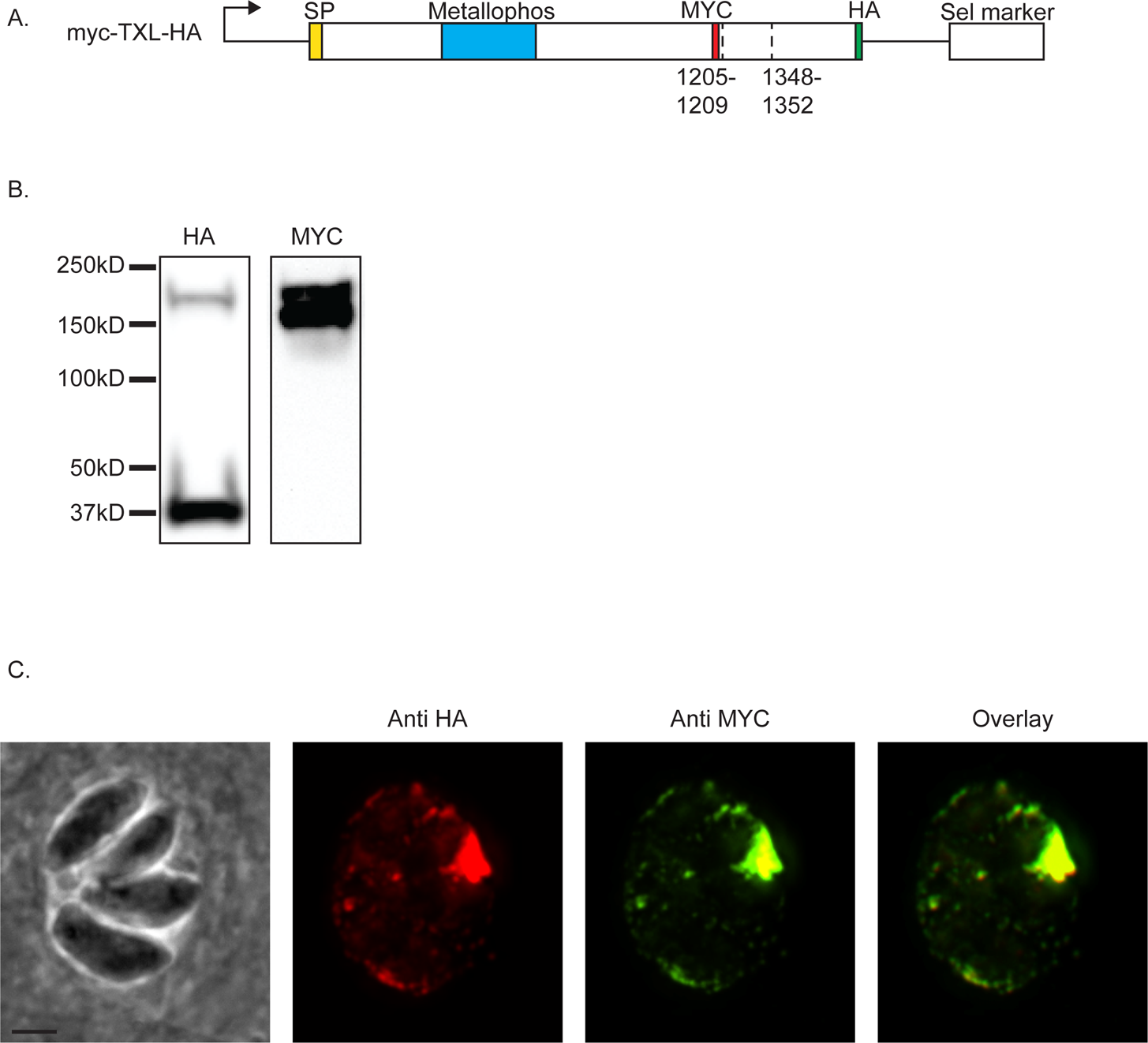
The GRA44 N-terminal cleavage product is secreted. To determine the stability and localization of the GRA44 N-terminal cleavage product we expressed an exogenous copy of GRA44 with an internal MYC epitope tag and a C-terminal HA epitope tag. A) Schematic of exogenous Gra44 construct myc-TXL-HA showing the position of the MYC and HA epitope tags relative to the two putative cleavage sites. B) Western blot of myc-TXL-HA parasite line separately probed anti-HA and anti-myc antibodies. C) IFA images of myc-TXL-HA expressing parasites probed for HA (red) and MYC (green). Scale bar = 2 µm.

### Loss of GRA44 negatively affects parasite propagation

Through a genome wide CRISPR screen, GRA44 was assigned a log_2_ relative fitness score of −3.28 (18), which would suggest that loss of GRA44 would be a significant detriment to parasite propagation. Accordingly, we applied a tetracycline (tet) repressible system (23, 24) to establish a conditional GRA44 knockdown strain. Specifically, we used CRISPR to introduce a cassette encoding a drug selective marker, a transactivator (TATi) protein and a tetracycline response element (TRE) just upstream of the endogenous GRA44 start codon (Fig. 4A). As to be able to monitor GRA44 protein expression we engineered the conditional knockdown mutant using the GRA44(HA) strain. The resulting strain, TATi-GRA44(HA) was grown for 24 and 48 hours in the absence and presence of the tetracycline analog anhydrotetracycline (ATc) and GRA44 expression was monitored by Western blot (Fig. 4B) and IFA (Fig. 4C). At both time points, protein levels are significantly reduced in presence of ATc when observed by either Western blot or IFA (Fig. 4B and C). Thus, we successfully established a strain in which expression of GRA44 can be conditionally turned down.

**Figure 4.**
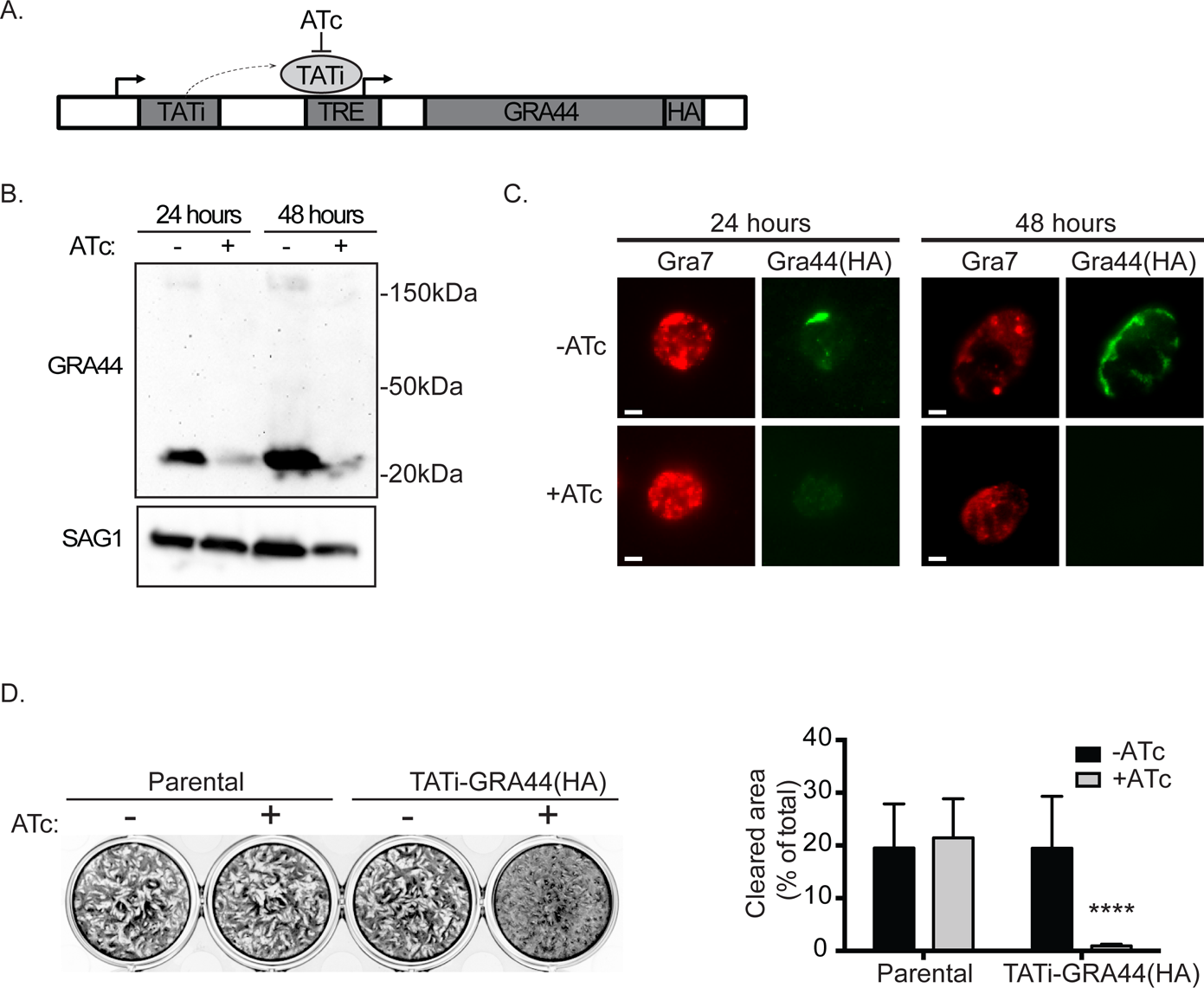
Knockdown of GRA44 disrupts parasite propagation. To determine the function of GRA44 we applied a tetracycline (tet) repressible system to establish a conditional GRA44 knockdown strain. A) Diagram of strategy used to generate conditional knockdown of GRA44, as outlined in methods section. B) Quantitative western blot of TATi-GRA44(HA) parasite strain grown for either 24 or 48 hours in absence (-) or presence (+) of ATc probed with HA to detect GRA44 and for SAG1 as a loading control. C) Reduction in GRA44 expression in presence of ATc was confirmed with IFA of intracellular parasites of the TATi-GRA44(HA) strain grown with and without ATc for 24 or 48 hours and probed with anti-HA antibodies. Scale bars = 2 µm. D) Plaque assays were performed with the GRA44(HA) parasites (parental) or the TATi-GRA44(HA) parasites grown without (-) or with (+) ATc for 5 days. Representative plaque assay is shown on the left. Results were quantitated based on percent cell monolayer cleared by parasite (cleared area) and the average biological and experimental triplicates are shown in graph (n=3, ±SD, p<0.0001 unpaired t test).

Interestingly, we noted the small C-terminal fragment from the TATi-GRA44(HA) strain was smaller than what is observed with the parental endogenous HA strain (Fig. 1B and 4B). Sequencing of GRA44 in the TATi-GRA44(HA) strain showed that a 315 base pair fragment, which encodes the last 105 amino acids, was deleted leaving the HA tag in frame. Surprisingly, we did not note any significant growth defect and successfully complemented with a full-length gene construct (Fig 4D, comparing parental to knockdown strain grown without ATc). Thus, deletion of that region does not affect localization or function. Regardless, this strain allows us to study the consequence of eliminating GRA44 expression upon ATc addition. In addition, to ensure that any phenotype is due to downregulation of GRA44 we complemented this strain with the addition of a wildtype copy of the gene (see below). Importantly, when the TATi-GRA44(HA) strain is grown in the presence of ATc, depleting GRA44, propagation is significantly affected as compared to the growth by the same strain under normal conditions (Fig. 4D). For the conditional knockdown strain, in the absence of ATc, we quantitated an average cell clearance of 19.4±9.9%, which was reduced to 0.9±0.4% when ATc was included in the growth medium (Fig. 4D). Therefore, conditional knockdown of GRA44 significantly reduces parasite propagation in tissue culture.

### Processing is not necessary for GRA44 function

To confirm the propagation defect observed upon knockdown of GRA44 was due to the reduction of GRA44 levels, we tested whether an exogenous copy of GRA44 could complement the phenotype. For this purpose, we introduced a copy of GRA44 with a C-terminal MYC epitope tag into the TATi-GRA44(HA) strain to generate a complemented strain TATi-GRA44(HA)comp, (Fig. 5A). The exogenous copy of GRA44(MYC) is processed and secreted as expected (Fig. 5B and C). In absence of ATc, both the endogenous HA-tagged GRA44 and the exogenous MYC-tagged GRA44 were detected in this strain by both Western blot and IFA (Fig. 5B and C). As expected, addition of ATc resulted in knockdown of GRA44(HA) but not of the exogenous GRA44(MYC) (Fig. 5B and C). Plaque assay of both the knockdown (TATi-GRA44(HA)) and complemented (TATi-GRA44(HA)comp) strains with and without ATc were performed in parallel to determine the ability of GRA44 to complement the phenotype. Consistent with the previous result, addition of ATc to the TATi-GRA44(HA) strain severely impaired plaque formation (Fig. 5D). Importantly, presence of a constitutively expressed copy of GRA44 complements this phenotype (Fig. 5D). While the percentage clearance of host cell in the presence of ATc was 0.8±0.4% for the knockdown strain after five days in culture, it was 26.6±3.7% for the complemented strain, which is statistically equal to what is observed without ATc with either strain (Fig. 5D). Complementation of the plaquing phenotype by the addition of a wildtype copy confirms that GRA44 is critical for efficient propagation of *Toxoplasma*.

**Figure 5.**
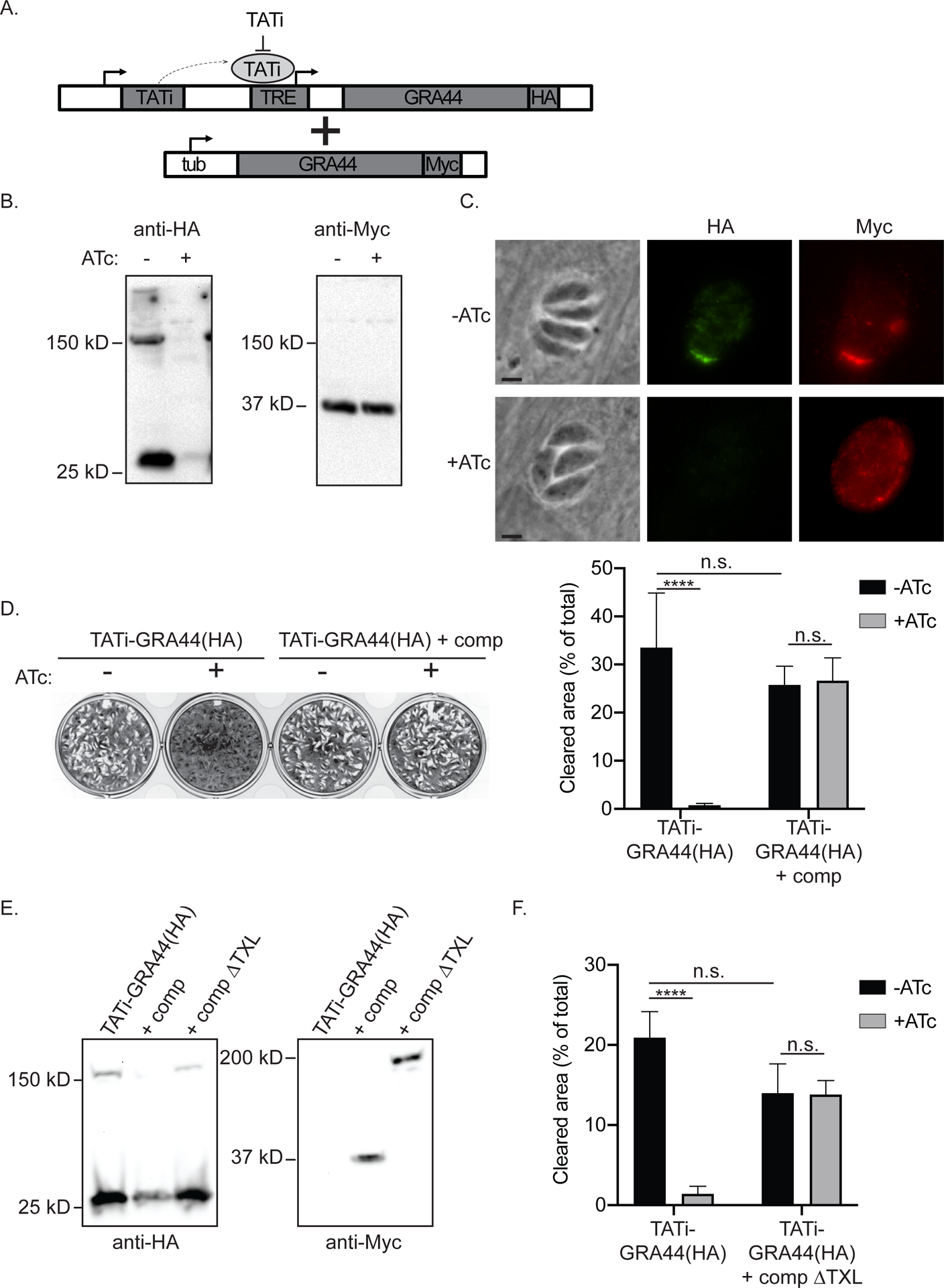
Phenotype of GRA44 knockdown can be complemented. To establish a direct connection between the lack of GRA44 and the phenotype observed, we established a complemented strain by adding an exogenous copy of wild type GRA44 to the TATi-GRA44(HA) conditional knockdown strain. A) Diagram of strategy used to establish a complemented strain TATi-GRA44(HA)comp. The wild type copy of GRA44 added to the TATi-GRA44(HA) strain contains a C-terminal MYC epitope tag and is driven by the *Toxoplasma* tubulin promoter (tub). B) Western blot of lysates from the TATi-GRA44(HA)comp parasites grown with and without ATc for 48 hours. Blots were probed for HA and MYC (C) Representative IFA images of TATi-GRA44(HA)comp with and without ATc treatment. In green is the HA tagged regulatable GRA44 and in red is the myc tagged constitutive exogenous copy. Scale bars = 2 µm. D) Plaque assays were performed with the knockdown (TATi-GRA44(HA)) and the complemented (TATi-GRA44(HA)comp) strains grown without (-) or with (+) ATc for 5 days. Average of biological and experimental triplicates is shown in bar graph based on percent of cell monolayer cleared (n = 3, ±SD, ****p<0.0001 One-way ANOVA followed by Tukey). E) TATi-GRA44(HA) was complemented with an exogenous copy of GRA44 containing TEXEL deletion 1348-1352 and C-terminal MYC tag (TATi-GRA44(HA)compΔTXL). Lysates from TATi-GRA44(HA), WT complemented strain TATi-GRA44(HA)comp and ΔTXL complemented strain TATi-GRA44(HA)compΔTXL were analyzed by Western blot and probed for HA and MYC. F) Plaque assays were performed with TATi-GRA44(HA) and TATi-GRA44(HA)compΔTXL strains. Parasites were grown 5 days without (-) or with (+) ATc. Average percent cell monolayer cleared for biological and experimental triplicates is shown by bar graph (n = 3 ±SD, **** p<0.0001 One-way ANOVA followed by Tukey).

Having established complementation of the growth phenotype, we set to determine whether processing of GRA44 was needed for function. Accordingly, we complemented the TATi-GRA44(HA) strain with an exogenous copy of GRA44(MYC) containing a TEXEL deletion of residues 1348-1352 to obtain the strainGRA44compΔTXL. For the GRA44compΔTXL strain under normal growth conditions without ATc, endogenous GRA44(HA) is detected by Western blot at a similar size to TATi-GRA44 and TATi-GRA44comp strains, however the exogenous MYC-tagged GRA44compΔTXL copy is seen as a mostly uncleaved form of the protein, in contrast to the wild type complement which is processed (Fig. 5E). Remarkably, the GRA44compΔTXL complemented parasite strain was no longer sensitive to the presence of ATc (Fig. 5E). These results indicate that complete cleavage of GRA44 is not necessary for function.

### GRA44 interacts with members of the effector translocation complex

Our results have thus far shown GRA44 to be of significant importance for successful parasite propagation, however the specific function of this protein within the parasitophorous vacuole is unclear. To shed light on its function we examined what proteins interact with GRA44 by developing a comprehensive interactome. For this purpose, GRA44 protein was immunoprecipitated from the GRA44(HA) tagged line and co-precipitating peptides analyzed by mass spectrometry. After three replicate experiments and controls with non-specific beads, the data were statistically analyzed by SAINT (Significant Analysis of Interactome) (25) computational predictive analysis (supplemental data set 1). With a SAINT score of >0.8 used as a cutoff, we obtained a list of 35 putative interactors, of which 8 are ribosomal and snRNP proteins and likely to be non-specific (Table S2 in supplemental material). Significantly, of the remaining 27 putative interactors, 23 have predicted signal peptides, which indicates that they are likely secreted proteins (Table 1). Among these are eight known GRA proteins (GRA9, 16, 25, 33, 34, 45, 50, and 52), the parasitophorous vacuole membrane associated protein MAF1, and MYR1 a known member of the effector translocation complex (26, 27).

**Table 1.** Putative GRA44 interactors identified by IP. Criteria used were Saint Score (SP) of >0.8, and not ribosomal protein (TGGT1_309820, TGGT1_207840, TGGT1_266070, TGGT1_248480). Highlighted proteins have a predicted signal peptide based on SignalP analysis. Proteins are listed based on SAINT score, total peptides and fold change over controls (table S2).

| ID number | Product description |
| --- | --- |
| TGGT1_316250 | GRA45 |
| TGGT1_204340 | GRA54 |
| TGGT1_254470 | MYR1 |
| TGGT1_319340 | GRA52 |
| TGGT1_279100 | MAF1a |
| TGGT1_251540 | GRA9 |
| TGGT1_203600 | GRA50 |
| TGGT1_304955 | serine/threonine specific protein phosphatase (PPM11C) |
| TGGT1_315610 | hypothetical protein |
| TGGT1_203290 | GRA34 |
| TGGT1_270320 | protein phosphatase 2C domain-containing protein (PPM3C) |
| TGGT1_258870 | hypothetical protein |
| TGGT1_311720 | chaperonin protein BiP |
| TGGT1_226240 | putative bud site selection protein |
| TGGT1_216770 | hypothetical protein |
| TGGT1_220950 | MAF1b |
| TGGT1_270240 | MAG1 |
| TGGT1_200360 | hypothetical protein |
| TGGT1_290700 | Gra25 |
| TGGT1_258458 | hypothetical protein |
| TGGT1_262050 | rhophry kinase family protein ROP39 |
| TGGT1_410360 | MAF1 copy |
| TGGT1_247440 | GRA33 |
| TGGT1_229480 | putative calcium binding protein precursor |
| TGGT1_208830 | GRA16 |
| TGGT1_410370 | MAF1 copy |

To confirm the interaction with MYR1, for which there are antibodies (26), we performed co-immunoprecipitating assays. Purified lysate from GRA44(HA) parasites was precipitated on mouse anti-HA beads, and eluates evaluated by Western for presence of GRA44 and MYR1. For MYR1, which is also processed by ASP5 at a TEXEL site, we probed with antibodies for either the C-terminal or N-terminal cleavage product. Immunoprecipitated GRA44 yielded a significant amount of either MYR1 fragment as compared to control IgG bead eluate of the same source (Fig. 6). Thus, GRA44 appears to interact with a member of the effector translocation system.

**Figure 6.**
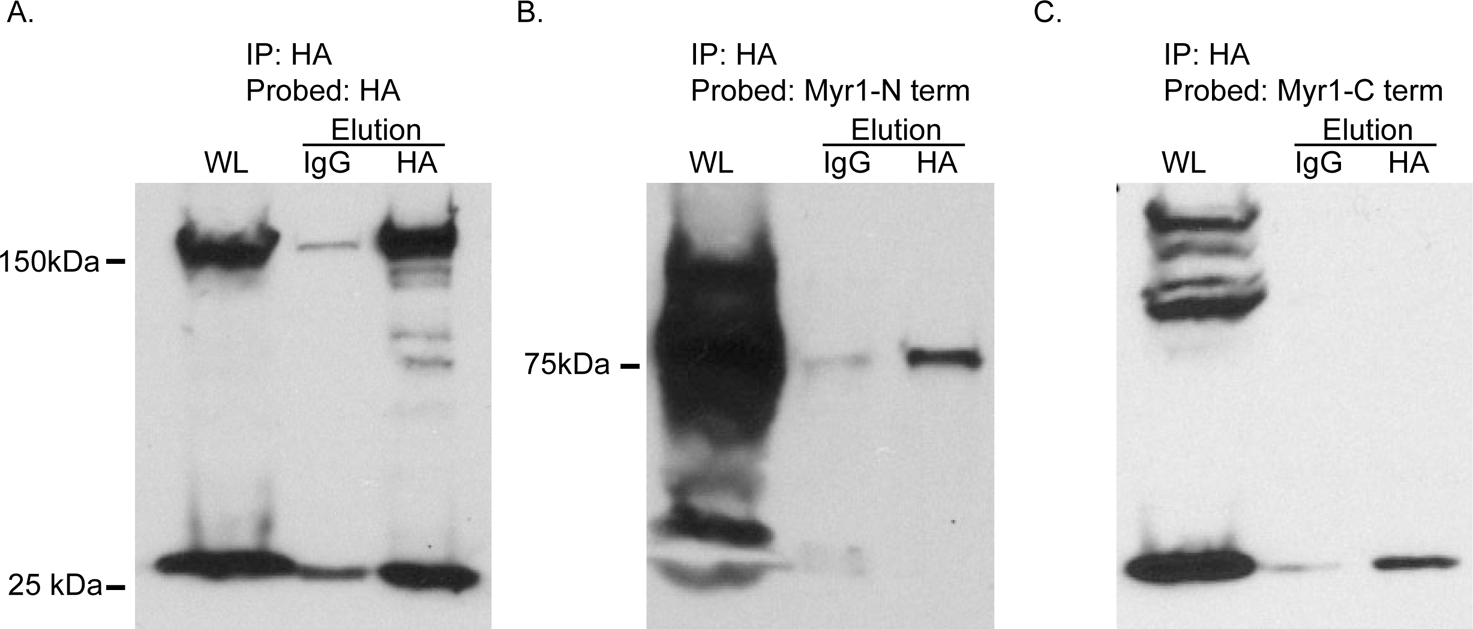
GRA44 interacts with MYR1. To confirm the interaction between GRA44 and MYR1, we performed immunoprecipitation from GRA44(HA) expressing parasites and probed for GRA44(HA) (A) or either the N-terminus (B) or the C-terminus (C) of MYR1 by Western blot. As controls we performed the immunoprecipitation was performed with IgG conjugated beads. In all blots the first lane is whole lysate (WL), middle lane is eluate from IgG beads, and last lane is eluate from the HA beads.

### GRA44 is required for cMYC induction

MYR1 was identified through a forward genetic screen to be required for translocation of parasite effectors such as GRA16 and GRA24 and the ensuing upregulation of the host cell oncogene cMYC (26). Given GRA44’s interaction with MYR1, we tested whether it might be involved in the same functions, specifically cMYC induction. For this purpose, TATi-GRA44(HA) parasites were grown for 24 hours either with or without ATc, released from host cells and allowed to infect new cells. Those that came from the +ATc conditions were kept in ATc, while those from the -ATc culture were kept without it. After 12 hours of growth, cultures were fixed and an IFA for human cMYC was performed (Fig 7). Images of merged phase contrast microscopy, HA, MYC, DAPi and cMYC staining were used to locate host cells infected by single PVs of greater than 1 parasite per vacuole and single channel images of cMYC staining were quantitated within these host nuclei boundaries by ImageJ software. Addition of ATc to TATi-GRA44(HA) parasites produced a reduction significantly reduced the level of GRA44 (Fig. 7A) detected consistent with the results observed by western blot and importantly reduced the cMYC signal by approximately 5-fold, a significantly dampened response as compared to normal conditions with the same strain (Fig. 7B). No significant difference in cMYC signal was observed in the complemented strain upon ATc addition.

**Figure 7.**
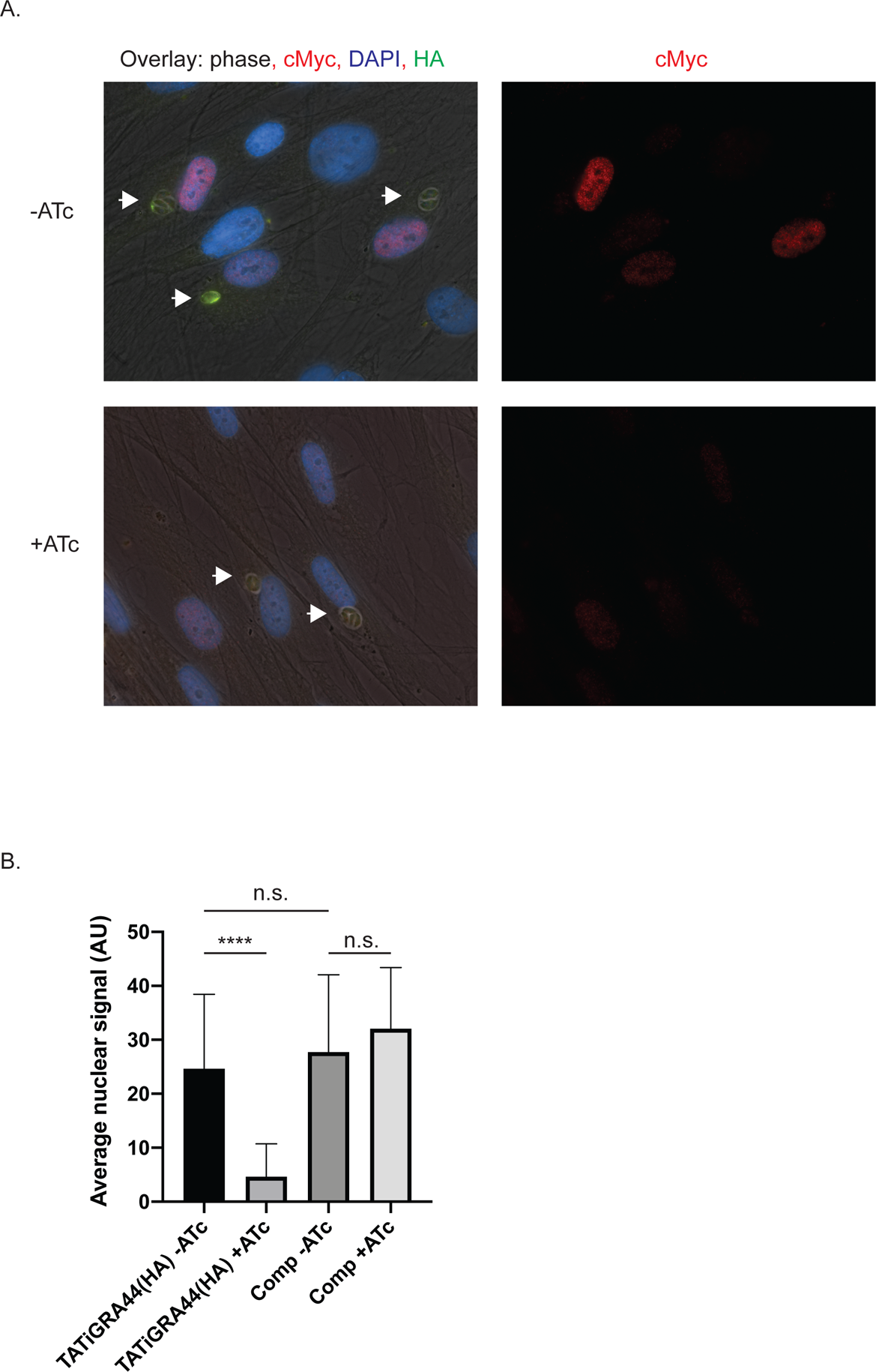
cMYC activation in GRA44 mutant strain. To assess host cell response upon GRA44 knockdown, host nuclear cMYC expression was quantified by fluorescence microscopy after invasion by either knockdown TATi-GRA44(HA) or complemented knockdown TATi-GRA44(HA)comp parasite strains in presence and absence of ATc. A. Representative immunofluorescence images of TATI-GRA44(HA) parasites grown without (-) or with (+) ATc Cultures were stained for host cMYC (red), DAPI to detect DNA and nuclei (blue), and HA to detect GRA44 (green). Arrows point at vacuoles with more than 2 parasites. B. Graph represents average quantified nuclear signal from cMYC antibody staining across biological and experimental triplicates. Arbitrary unites (AU) were used in comparing nuclear cMYC signal intensity. (n= 3, ****p<0.0001 One-way ANOVA followed by Tukey).

## DISCUSSION

As part of its lifecycle, *Toxoplasma* invades and dwells within host cells where it utilizes nutrient resources readily available within a host. Since residence inside a host is key to parasite survival and propagation, *Toxoplasma* parasites have multiple means of altering the status of their host to better suit their needs. Examples of changes induced within the host include apoptosis inhibition (14), innate immune system disruption (28), host cytoskeleton restructuring (29) and global changes to host gene transcription (9). These critical host cell alterations are accomplished by a large arsenal of parasite effectors that are secreted into the host cell during invasion and intracellular growth. Secretion of these effectors involve highly coordinated actions by two unique parasite organelles: the rhoptries and the dense granules. The rhoptries discharge proteins known as ROPs during invasion, while the dense granules release their contents into the PV during intracellular growth. While some of these so called GRA proteins remain within the PV space or associate with the PV membrane (PVM), many of them are translocated across the PVM into the host where they affect numerous signaling pathways. How these proteins move from the PV to the host is becoming clear with the discovery of proteins that appear to form part of a putative translocon (24). The work presented here shows that GRA44 (TGGT1_228170), previously known as IMC2A (16), interacts with members of this complex and that it is essential for some of the host cell events downstream of effector translocation.

Work from Coffey et al. identified GRA44 as a substrate of ASP5 through a comparative proteomic approach (13). They showed that indeed GRA44 is processed and secreted into the PV, which we have corroborated with the work presented here. Their analysis of GRA44 identified two cleavage sites, one at 83-85 and another at 1348-1350. Nonetheless, their studies did not determine whether all cleavage products were secreted or the function/role of GRA44 within the PV. Here we report that the two major GRA44 products are secreted into the PV. This is important as it is the first evidence that the fragment containing the putative acid phosphatase is indeed in the PV. In our study, we show that the processing is not necessary for either the secretion or function of GRA44. This is consistent with the fact that other secreted proteins containing TEXEL motifs such as MYR1 and WNG1/2 can still be secreted in absence of ASP5 activity (13). Fortuitously, through a spontaneous deletion in the knockdown strain, we also determined that the last 105 amino acids of the protein are not needed for function or localization.

Proteomic analysis of proteins co-immunoprecipitated with GRA44, reveal interaction with known secreted proteins, including numerous GRAs. Among these interactors are GRA9, part of the intravacuolar tubular network (30), GRA16 an effector altering host cell cycle through p53 pathway (31), GRA25 a macrophage-dependent immune modulator (32), GRA33, GRA34, GRA45, GRA50, GRA52 and members of the multi-copy Mitochondrion Association Factor 1 (MAF1) family. MAF1 is involved in recruiting the host mitochondria to the PV membrane surface (33). Most importantly, immunoprecipitation revealed an interaction between MYR1 and GRA44, which we have confirmed independently by Western blotting (Fig. 6). Interestingly, as we were studying the interactome of GRA44 we learned of ongoing work in Dr. Boothroyd’s lab showing a physical and functional interaction between MYR1 and GRA44 (see accompanying manuscript). MYR1 was initially identified through a forward genetic screen for *Toxoplasma* mutants unable to activate host cMYC and translocate effectors such as GRA16 and GRA24 (22). Further screening for such mutants identified MYR2 and MYR3 as also required for effector translocation (27). Thus, MYR1/2/3 appear to be part of a putative translocon complex, although only MYR1 and MYR3 have been confirmed to directly interact (27). MYR1-dependent effectors are responsible for driving a broad range of host cell effects early on during infection. MYR1-dependent host responses include upregulation of E2F transcription factors and downregulation of interferon signaling (8). The interaction between GRA44 and MYR1 would suggest a function in translocation for this putative acid phosphatase.

One of the challenges in determining the relevance of interactions among dense granule proteins is that those interactions could be within the dense granules or during transit and not necessarily once in the PV or host where they exert their function. Nonetheless, our data strongly suggest that the interaction with MYR1 is functionally relevant. Similar to knockout of MYR1, knockdown of GRA44 results in significant reduction of cMYC activation. Together, these findings support the notion that GRA44 is a member of the MYR1/2/3 translocon machinery. Whether its role in this process is structural or regulatory remains unknown and would require further investigation. The possibility that GRA44 plays a regulatory function is suggested by the presence of a putative catalytic site reminiscent of acid phosphatases. Whether GRA44 is an active phosphatase or a pseudophosphatase remains to be determined.

In the biology of animals, plants and fungi, acid phosphatases serve many biological purposes and proteins are designated as such based on a shared similarity in catalytic site structural arrangement (34). Typically these enzymes coordinate an Fe(III) and divalent metal such as Mn(II), Zn(II) or Fe(II) as part of their active site. Each metal is coordinated to three amino acids and shares a linking aspartic acid bridge between them with the Fe(III) typically mated to a histidine, an asparagine and a tyrosine and the divalent metal bound by two histidines and an asparagine. GRA44 contains a majority of the conserved residues common to acid phosphatases. In GRA44 the aspartic acid conjugating the Fe(III) is switched to an asparagine and the asparagine conjugating the second metal is replaced by a glutamic acid. This would represent a swap in charged amino acids, and should still maintain a stable charge equilibrium within the active site. The only coordinating residue unaccounted for is the tyrosine binding Fe(III). The active site of GRA44 could then be hypothesized to consist of an Asp bridge between the first metal M(II), bound by a glutamic acid and two histidines, and the second metal M(III), bound by an asparagine, a histidine, and an unknown seventh residue.

Acid phosphatases commonly scavenge, recycle and transport inorganic phosphorous, and have been implicated in various biological functions, such as downregulation of prostate cell growth signaling and osteoclast bone resorption activity (35, 36). Acid phosphatases have also been implicated in phosphate acquisition from organophosphate compounds and dephosphorylative regulation of enzymes in plants (37, 38). GRA44 function, at least in part, is likely related to its interactions with the MYR translocon in the PV. Two plausible functions for GRA44 could be regulation of MYR component proteins by dephosphorylation or dephosphorylation of effectors for trafficking across the PVM structure. Typically, proteins must be dephosphorylated to cross a lipid bilayer membrane such as the PVM and notably both GRA16 and GRA24 have been shown to be phosphorylated as identified by *Toxoplasma* phosphoproteome analysis (39)). MYR3 also exists in a partial phosphorylated state and could be a substrate for a phosphatase such as GRA44 (27). Consistent with the idea of phosphoregulation of the export system, the secreted kinase ROP17 has been shown to be critical for efficient effector translocation (40).

Besides the defect in cMYC activation, genetic disruption of GRA44 results in a propagation defect. This result is congruent with published work describing the effect of complete GRA44 knockout (13) and with the fitness score of −3.28 assigned to GRA44 through a genome wide CRISPR screen (18). This is particularly interesting, as disruption of other translocon members do not affect fitness to the level seen with GRA44. For example, the relative fitness scores for MYR1, MYR2, MYR3 are 0.88, 2.39 and 2.83 respectively, although disruption of any of these interferes with effector translocation and cMYC activation. Similarly, disruption of the effectors GRA16, GRA24 and TgIST, which depend on MYR1 for translocation, have positive fitness scores of 1.44, 2.28 and 2.86. Thus, it is unlikely that the growth defect exhibited by parasites lacking GRA44 is due to defects on effector translocation. Alternatively, GRA44 might play several independent roles, including nutrient acquisition, which would be consistent with known functions of acid phosphatases.

In conclusion, we have shown GRA44 to be a secreted protein critical for *Toxoplasma* survival and propagation that plays a significant role in host manipulation and interacts with the translocon protein MYR1. As part of its secretion to the PV it is cleaved at an internal TEXEL site forming two stable and colocalizing proteins. The mechanistic action by which GRA44 is involved with protein secretion to host cells remains unknown, however due to its putative acid phosphatase domain, involvement with dephosphorylation of trafficked proteins or members of the translocon complex is plausible. Future work on the activity and substrates of GRA44 will shed light on the regulation of effector translocation, a process central to the interactions between *Toxoplasma* and its host.

## MATERIALS AND METHODS

### Parasite and Host Cell Culture

All parasite lines were maintained by continuous passage through human foreskin fibroblasts (HFFs) purchased from ATCC. Parasites and HFFs were grown in Dulbeco’s Modified Eagle Medium (DMEM) supplemented with 10% fetal bovine serum (FBS), 2mM/L glutamine and 100 units penicillin/100μg streptomycin/ml. When pyrimethamine was included in media for selection, dialyzed FBS was used. All parasite and HFF cultures were grown in a humidified incubator at 37°C with 5% CO_2_. Initial parental parasite lines used were RH strain lacking hypoxanthine-xanthine-guanine phosphoribosyl transferase (HPT) gene, referred to as RHΔ*hpt* (41), and RH strain lacking HPT and Ku80, referred to as RHΔ*ku80* (42, 43). For drug treatment and selection, stocks of pyrimethamine and chloramphenicol were prepared in ethanol, stocks of anhydrotetracycline (ATc) were prepared in DMSO. All drugs were purchased from Sigma.

### Endogenous epitope tagging

For C-terminal endogenous tagging of TGGT1_228170, the 3’ region directly upstream of the stop codon was amplified from RhΔ*ku80* parasite genomic DNA by PCR and inserted into the pLIC-3xHA-DHFR (43) vector at the PacI restriction site by ligation independent cloning (LIC) facilitated by InFusion HD Cloning Plus (Clontech). Sequences of primer for this and all reactions used in this work are in supplemental table S3. 50µg of XcmI linearized vector was transfected into RhΔ*ku80* parasites and the resultant population was selected for the presence of the pyrimethamine resistant dihydrofolate reductase (DHFR) allele, which is included in the vector (44). Independent clones were established by limiting dilution of the transfected population and confirmed by immunofluorescence assay and Western blots.

### Exogenous gene insertion and parasite line generation

To introduce an exogenous copy of TGGT1_228170 into parasites, we first generated a vector containing a section of the genomic TGGT1_228170 locus beginning from the start codon to the stop codon that included introns and a C-terminal HA epitope. The section of TGGT1_228170 in the vector was flanked by the *Toxoplasma* tubulin promoter and 5’UTR and the tubulin 3’UTR. This was achieved by cloning a PCR amplicon of the TGGT1_228170 genomic DNA (primers in table S3) into the NcoI and PacI sites of pTNRluc-Tub-HPT (45) using InFusion HD Plus for LIC. 50µg of the resulting vector, pTub-Gra44-HPT, linearized with ScaI, was transfected into RhΔ*hpt* parasites. The transfected population was selected for HPT by adding mycophenolic acid (50μg/mL) and xanthine (50μg/mL) to the media. Independent clones were established by limiting dilution. For mutant variations of the exogenously expressed TGGT1_228170, TEXEL deletions, myc epitope tag insertion and TEXEL2 point mutations were introduced into pTub-Gra44-HPT using the Q5 site directed mutagenesis kit (NEB) and TEXEL point mutations were accomplished similarly with Quikchange site-directed mutagenesis kit (Agilent).

### Development of GRA44 conditional knockdown parasite line

To generate the GRA44 conditional knockout strain we introduced a cassette encoding a drug selective marker, a transactivator (TATi) protein and a tet response element (TRE) just upstream of the GRA44 start codon. This TATi cassette was amplified from the vector pT8TATi-Gra44-HX-tetO7S (46) with primers that include areas of homology upstream of GRA44 to facilitate homologous recombination. 1μg of this PCR amplicon was transfected into the RhΔKu80 parasites that express endogenously HA tagged GRA44. To drive the insertion of the TATi cassette we co-transfected the PCR amplicon with 2μg of a vector expressing Cas9 and a guide RNA targeting the TGGT1_228170 locus upstream of the start codon. This vector was made using pSAG1-Cas9-GFP-pU6-sgUPRT (47) as a template and the sequences encoding the guide RNA were introduced with Q5 site directed mutagenesis (NEB). Parasites transfected with the TATi cassette and Cas9 vector were selected for HPT and independent clones established by limiting dilution. Correct integration of the TATi insert cassette was validated by PCR. Resulting strain was designated TATi-GRA44(HA).

### Complementation of conditional knockdown line

To complement the knockdown strain, a wildtype copy of TGGT1_228170 driven by a *Toxoplasma* tubulin promoter and including a C-terminal MYC epitope tag was targeted to the inactive Ku80 locus of theTATi-GRA44(HA) strain using CRISPR/Cas9 to assist integration. The insertional cassette, which includes the tubulin-driven TGGT1_228170 and a chloramphenicol resistance gene (48) was amplified by PCR from plasmid pTub-Gra44-myc-CmR with primers that included homology segments to the Ku80 locus (table S3).The pTub-Gra44-myc-CmR vector was constructed with Infusion assisted LIC cloning by inserting the Gra44 gene with appended C-terminal myc tag, amplified from pTub-Gra44-HPT, to the pLIC-SMGFP-CmR vector backbone (49) replacing the SMGFP tag and upstream region. 1µg of this PCR amplicon was co-transfected with 2 µg of Cas9 vector encoding an sgRNA targeting the Ku80 locus.

### SDS-PAGE and Western blot analysis

For detection of protein in lysates from extracellular parasite samples, parasites were allowed to undergo natural egress then collected, centrifuged and washed 2x with cold PBS (10 min, 1,000 x g). For analysis of intracellular parasite protein lysates, host cell monolayers were washed 2x with cold PBS, scraped and centrifuged for 10 minutes at 1,000 x g. Parasite samples were resuspended in 2X sample loading buffer with 5% β-mercaptoethanol and boiled for 5 minutes at 98°C. Boiled samples were frozen at −20°C, then thawed and re-boiled for 5 minutes at 98°C before gel loading. SDS-PAGE and Western blots were performed with standard methods as previously described (50).

For Western blot analysis of GRA44 conditional mutant strains, parasites were first grown under normal conditions for 24 hours, then syringe lysed with a 27-gauge needle. Fresh host cells were infected with an equal quantity of syringe lysed parasites and grown for 24 or 48 hours with or without 1μg/mL ATc. For analysis of protein lysates from extracellular parasite samples, host cells were scraped and parasites released by passing through a syringe and centrifuged 10 minutes at 1,000 x g. For analysis of intracellular parasite protein lysates, host cell monolayers were washed with cold PBS, scraped and centrifuged for 10 minutes at 1,000 x g. Resulting samples were resuspended in 200 μL RIPA lysis buffer (50mM Tris, 150 mM NaCl, 0.1% SDS, 0.5% sodium deoxycholate, 1% TritonX-100) including protease/phosphatase inhibitor cocktail (Cell Signaling Technology) and incubated on ice 1 hour, sonicated 2 times for 15 seconds with 1 minute rests on ice and centrifuged (20,000 x g, 15 minutes, 4°C). Supernatants were combined with 4X SDS loading buffer with 10% β-mercaptoethanol and boiled for 5 minutes at 98°C. Boiled SDS samples were frozen at −20°C, then thawed and re-boiled for 5 minutes at 98°C before gel loading. SDS-PAGE and Western blots were performed with standard methods as described above. Uncropped original images for all western blots are included as a supplemental data set 2.

Primary antibodies used for western blots included rabbit anti-HA at a dilution of 1:1,000 (Cell Signaling Technologies), rabbit anti-MYC at a dilution of 1:1,000 (Cell Signaling Technologies), mouse anti-SAG1 at a dilution of 1:2,000 (Genway), and mouse anti-Myr1 antibodies at 1:1,000 (26, 27). Secondary antibodies used include peroxidase-conjugated goat anti-mouse and anti-rabbit and were used at a 1:10,000 dilution.

### Immunofluorescence assays

For all immunofluorescence assays (IFA), HFFs were grown to confluency on 1.5mm glass coverslips and infected with parasites, which were allowed to grow for 20 hours prior to fixation with 4% paraformaldehyde for 20 minutes. Cells were washed 1x with PBS after fixation, then permeabilized and blocked with a solution of 3%BSA/0.2%Triton-X100 in PBS for 15-20 minutes. Coverslips were incubated with primary antibodies in 3%BSA/0.2%Triton-X100 in PBS for one hour at room temperature and washed five times with PBS. Finally, cultures were incubated with fluorophore-conjugated secondary antibodies for one hour at room temperature in 3%BSA in PBS, then washed five times with PBS and mounted on glass slides with vectashield mounting medium containing DAPI (Vector Laboratories). Primary antibodies used were rabbit anti-HA at 1:1000, mouse anti-MYC at 1:1000 (Cell signaling Technology), rat anti-HA at 1:2000 (Roche), rabbit anti-human C-myc at 1:1000 (Abcam), mouse anti-gra5 (Biotem) at 1:1000, and mouse anti-gra7 at 1:1000. Secondary antibodies used (Life Technologies) were Alexafluor-488 goat anti-rabbit or goat anti-rat, alexafluor-594 goat anti-mouse or Goat anti-Rabbit and Alexafluor-647 Goat anti-Mouse. All secondary antibodies were used at 1:2000. Images were taken on a Nikon Eclipse 80i microscope using a Nikon DS-Qi1Mc camera and NIS Elements AR 3.0 software.

### Immunoprecipitation and Co-IP experiments

Infected host cells were washed 2x with cold PBS and scraped from the flask surface to collect intracellular parasites, which were centrifuged 10 minutes at 1,000 x g and resuspended in 200 μL ice cold IP lysis buffer (Pierce, Thermo Scientific) containing protease and phosphatase inhibitors (Cell Signaling Technology). Lysate was incubated on ice one hour, sonicated on ice 2x for 15 seconds and centrifuged 15 minutes at 20,000 x g and 4°C. Supernatant was collected and incubated with magnetic beads conjugated to either mouse IgG or primary antibody (Pierce, Thermo Scientific) for one hour at 4°C with rocking. Incubated beads were separated from solution with a magnet and washed with IP lysis buffer (Pierce, Thermo) plus inhibitors three times and either stored in 8M Urea at −80°C for downstream mass spectrometric analysis or directly eluted into 2x SDS sample loading buffer/5% β-mercaptoethanol, boiled 5 minutes at 98°C, and stored at −20°C for Western blot analysis. SDS-PAGE and Western blots were performed as outlined above. Protein analysis by mass spectrometry was completed by Indiana University School of Medicine Proteomics Core facility as previously described (50).

### Plaque assays

12-well plates were infected with 500 parasites/well of freshly syringe-lysed parasites and grown undisturbed for 5 days before fixation with methanol for 5 minutes. Wells were stained with crystal violet and plaque images quantified by ImageJ using the ColonyArea plugin (50).

### HFF cMYC response assay and quantitation

Parasites used for C-myc assays were grown 48 hours with or without 1 μg/mL ATc and syringe-lysed prior to infection of host cell coverslips. Coverslips of confluent HFF monolayers pretreated for 24 hours with FBS-free media were infected and fixed 19 hours post infection. IFAs for human C-myc, HA and DAPI were performed as described above. Images of phase contrast, DAPI, HA, myc and C-myc channels were acquired for at least 20 vacuoles under each experimental condition and exported to ImageJ. Infected host cell nuclei were identified from merged-channel images and quantitated for C-myc expression from images of the C-myc channel alone. Measurements were taken of mean pixel intensity within host nucleus boundaries of singly infected cells containing PVs greater than one parasite. Measurements from triplicate experiments were averaged.

## ACKNOWLEDGMENTS

This work was funded by grants from the National Institutes of Health to GA (R01AI123457 and R21AI138255). We would like to thank Jason True and other members of the Indiana University School of Medicine Proteomics Core facility for mass spectometric sample analysis and data processing. In addition we would like to thank Alicja Cygan, Terence Theisen and Dr. John Boothroyd for collaborative discussions and sharing of reagents and unpublished data with regard to this work.

