## Supplemental material for "The secreted acid phosphatase domain-containing GRA44 from *Toxoplasma gondii* is required for C-myc induction in infected cells"

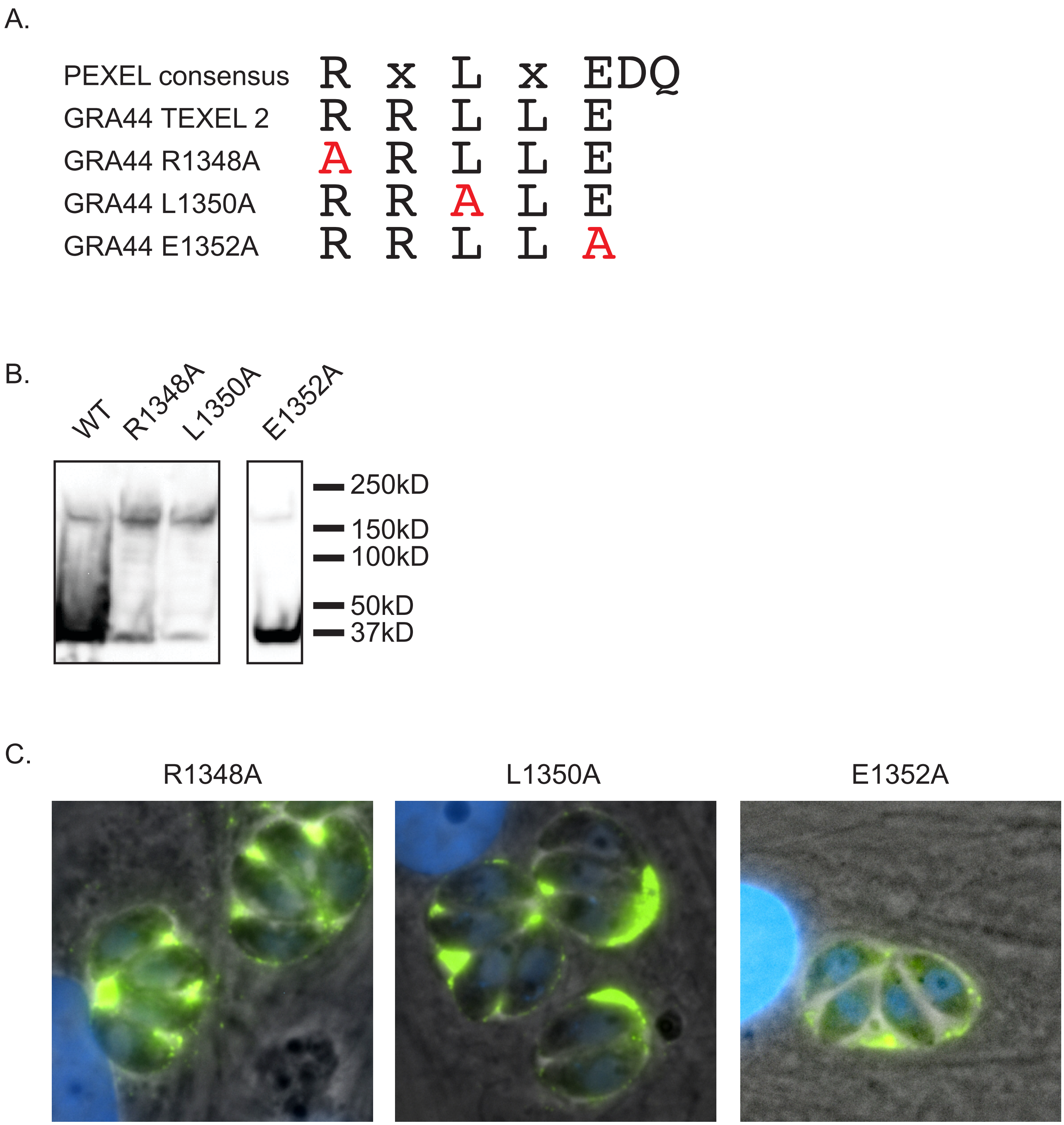


**Figure S1. Mutational analysis of TEXEL2.** A) Alignment of the PEXEL consensus, and the GRA44 TEXEL along with the three mutant versions R1348A, L1350A and E1352A. B) Western blot of lysates from parasites expressing exogenous wild type or mutant GRA44 probed with HA antibodies. D) Representative IFA images of intracellular parasites expressing each of the four mutant GRA44. Images are overlay of phase and HA signal (in green).

### Supplemental figure S2

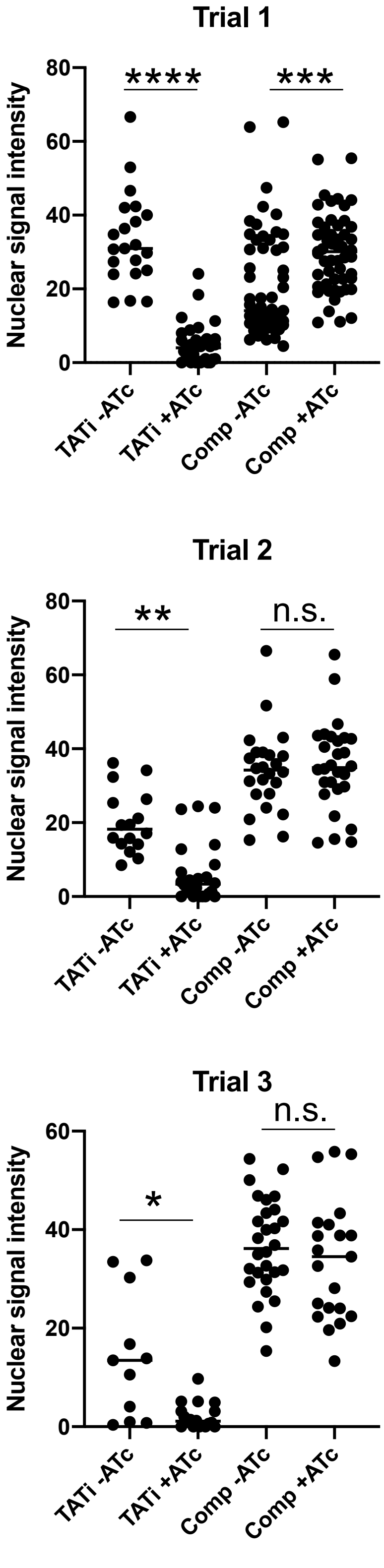


Figure S2. c-Myc activation by parasites of the TATi-GRA44(HA) and complemented strains grown with or without ATc. Host cells were monitored for nuclear c-Myc staining. Results from three independent experiments are shown. Data was analyzed by ANOVA, **** $p<0.0001$, *** $p=0.0002$, ** $p=0003$, * $p=0.01$

| ID number | Product Description | Fitness score |
|---|---|---|
| TGGT1_283720 | phosphotyrosyl phosphate activator (ptpa) protein | -4.71 |
| TGGT1_224920 | hypothetical protein | -4.56 |
| TGGT1_311290 | protein tyrosine phosphatase family protein, ptpla protein | -3.86 |
| TGGT1_283590A | NLI interacting factor family phosphatase | -3.76 |
| TGGT1_219320 | acid phosphatase GAP50 | -3.74 |
| TGGT1_228170 | inner membrane complex protein IMC2A | -3.28 |
| TGGT1_276210 | phosphoglycerate mutase family protein | -2.94 |
| TGGT1_216600 | exonuclease III APE | -2.67 |
| TGGT1_237410 | protein phosphatase 2C domain-containing protein | -2.01 |
| TGGT1_204080 | histidine acid phosphatase superfamily protein | -1.99 |
| TGGT1_243990 | Dullard family phosphatase domain-containing protein | -1.94 |
| TGGT1_224220 | serine/threonine-protein phosphatase PP2A catalytic subunit | -1.5 |
| TGGT1_305910 | hypothetical protein | -1.31 |
| TGGT1_252380 | hypothetical protein | -1.01 |
| TGGT1_259960 | Nucleoside-diphosphatase | -0.67 |
| TGGT1_204410 | endonuclease/exonuclease/phosphatase family protein | -0.49 |
| TGGT1_201630A | protein phosphatase 2C domain-containing protein | -0.06 |
| TGGT1_277720 | GDA1/CD39 (nucleoside phosphatase) family protein | 0.28 |
| TGGT1_244450 | protein phosphatase 2C domain-containing protein | 0.51 |
| TGGT1_228160 | acid phosphatase | 0.55 |
| TGGT1_278878 | GDA1/CD39 (nucleoside phosphatase) family protein | 0.6 |
| TGGT1_268770 | dual specificity phosphatase, catalytic domain-containing protein | 0.71 |
| TGGT1_278882 | GDA1/CD39 (nucleoside phosphatase) family protein | 0.71 |
| TGGT1_276920 | protein phosphatase 2C domain-containing protein | 0.75 |
| TGGT1_308950 | histidine acid phosphatase superfamily protein | 0.77 |
| TGGT1_225290 | GDA1/CD39 (nucleoside phosphatase) family protein | 0.81 |
| TGGT1_278510 | protein phosphatase 2C domain-containing protein | 1.01 |
| TGGT1_237500 | protein phosphatase 2C domain-containing protein | 1.13 |
| TGGT1_270320 | protein phosphatase 2C domain-containing protein | 1.24 |
| TGGT1_297650 | Ser/Thr phosphatase family protein | 1.52 |
| TGGT1_222840 | Ser/Thr phosphatase family protein | 2.25 |

Table S1. List of proteins annotated as phosphatases and predicted to contain a signal peptide. Fitness score is the mean phenotype score based on a genome wide CRISPR screen (15).

| ID number | Product description | total peptides IPs | Total peptides controls | Fold Change | SAINT Score |
|---|---|---|---|---|---|
| TGGT1_316250 | GRA45 | 111 | 0 | INF | 1 |
| TGGT1_262960 | putative U1 snRNP-associated protein Usp106 | 44 | 0 | INF | 1 |
| TGGT1_204340 | hypothetical protein | 39 | 0 | INF | 1 |
| TGGT1_254470 | MYR1 | 30 | 0 | INF | 1 |
| TGGT1_319340 | GRA52 | 27 | 0 | INF | 1 |
| TGGT1_279100 | MAF1 copy | 20 | 0 | INF | 1 |
| TGGT1_309820 | ribosomal protein RPL11 | 15 | 0 | INF | 1 |
| TGGT1_228170 | IMC2A /GRA44 | 1023 | 19 | 54 | 1 |
| TGGT1_251540 | GRA9 | 49 | 1 | 49 | 1 |
| TGGT1_203600 | GRA50 | 32 | 1 | 32 | 1 |
| TGGT1_304955 | serine/threonine specific protein phosphatase | 31 | 1 | 31 | 1 |
| TGGT1_207840 | ribosomal protein RPS17 | 47 | 2 | 24 | 1 |
| TGGT1_315610 | hypothetical protein | 21 | 1 | 21 | 1 |
| TGGT1_203290 | GRA34 | 13 | 1 | 13 | 1 |
| TGGT1_266070 | ribosomal protein RPL31 | 13 | 1 | 13 | 1 |
| TGGT1_270320 | protein phosphatase 2C domain-containing protein | 13 | 0 | INF | 0.99 |
| TGGT1_258870 | hypothetical protein | 33 | 2 | 17 | 0.99 |
| TGGT1_311720 | chaperonin protein BiP | 140 | 26 | 5.4 | 0.99 |
| TGGT1_226240 | putative bud site selection protein | 7 | 0 | INF | 0.97 |
| TGGT1_216770 | hypothetical protein | 6 | 0 | INF | 0.96 |
| TGGT1_242330 | ribosomal protein RPS5 | 60 | 12 | 5 | 0.95 |
| TGGT1_220950 | MAF1 copy | 19 | 2 | 9.5 | 0.94 |
| TGGT1_270240 | MAG1 | 112 | 35 | 3.2 | 0.94 |
| TGGT1_200360 | hypothetical protein | 24 | 3 | 8 | 0.92 |
| TGGT1_290700 | GRA25 | 15 | 1 | 15 | 0.91 |
| TGGT1_258458 | hypothetical protein | 10 | 1 | 10 | 0.89 |
| TGGT1_262050 | rhoptry kinase family protein ROP39 | 39 | 2 | 20 | 0.88 |
| TGGT1_410360 | MAF1 copy | 19 | 2 | 9.5 | 0.87 |
| TGGT1_248480 | ribosomal protein RPS9 | 32 | 5 | 6.4 | 0.86 |
| TGGT1_231140 | ribosomal protein RPS25 | 14 | 2 | 7 | 0.85 |
| TGGT1_247440 | GRA33 | 15 | 2 | 7.5 | 0.84 |
| TGGT1_229480 | putative calcium binding protein precursor | 26 | 4 | 6.5 | 0.84 |
| TGGT1_208830 | GRA16 | 14 | 2 | 7 | 0.83 |
| TGGT1_410370 | MAF1 copy | 23 | 4 | 5.7 | 0.8 |
| TGGT1_267400 | ribosomal protein RPL32 | 26 | 5 | 5.2 | 0.8 |

Table S2. Proteins identified through immunoprecipitation and MS/MS with SAINT score of 0.8 or higher. Number of peptides shown are total for all three experiments or controls and the fold change is number of peptides in experimental IPs over that of control ones.

| Primer Use | Primer # | Primer Name | Sequence |
|---|---|---|---|
| Cloning Gra44 C-terminal region into endogenous tagging vector | 1 | IMC2A-HA-LIC.For | ttccaatccaatttaattaaagacagcacgggaacttgc |
| | 2 | IMC2A-HA-LIC.Rev | ccacttccaattttaattaattccgttgcgtcagtcg |
| Inserting Gra44 gDNA to pTNRLUC-HX | 3 | IMC2A-TNRLUC.FOR | TTTCGACAAAccATGGAAAGACGTACACGTCCG |
| | 4 | IMC2A-TNRLUC.REV | CGGGCTTGCGGTttaattaaTTAGGCATAATCTGGAACATCGTAAGGATAttc cgttgcgtcagtcgagtc |
| Amplification of TATi cassette for insertion at Gra44 promoter | 5 | TATi-IMC2A.F | GCTCGGTATTCATACCTAGAGATTCCTCTGTTTTCGAACAAttctcatgtttgcg gatccg |
| | 6 | TATi-IMC2A.R | CCGGACGTGTACGTCTTTCCATCTTTGTAGAAACGTGGGCcaggtcctcctcg gagatga |
| Q5 insertion of Gra44 promoter sgRNA to Cas9 plasmid | 7 | TATi-IMC2A sgRNA.F | tcattgtcccGTTTTAGAGCTAGAAATAGC |
| | 8 | TATi-IMC2A sgRNA.R | tcgtccccgcAACTTGACATCCCCATTTAC |
| Insertion and tagging of Gra44 gene to complement vector | 9 | IMC2A-myc CmR.F | ACGGGAATTCCCTAGATTGGGTACCGGGCCC |
| | 10 | IMC2A-myc CmR.R | CAACTTTTCTACATATTACAGGTCCTCCTCGGAGATCAGCTTCTGCTCttc cgttgcgtcagtcgag |
| Amplification of Gra44 complement cassette for Ku80 site insertion | 11 | IMC2A-CmR [Ku80].F | GTCCCCGGTTCGCCTCAGCACACACACACATGACGTACATCGAAGCTG GGTACCCTGTACTTCC |
| | 12 | IMC2A-CmR [Ku80].R | GTAATGTCGGAATAGTTCCCATCAGAAACAATGGAGCTATCCGCGTCCC ATTCGCCATTCAG |
| Insertion of Ku80 sgRNA to Cas9 plasmid | 13 | sgKU80.F1 | ctcatattccGTTTTAGAGCTAGAAATAGC |
| | 14 | sgKU80.R1 | aaaggtgtacAACTTGACATCCCCATTTAC |
| Myc epitope insertion upstream of TEXEL1 | 15 | myc-PEXEL1.F | tccgaggaggacctgCGGAGAGAGCTAGAGGAAC |
| | 16 | myc-PEXEL1.R | gatcagcttctgctcCAGTCCGCCAATCGATCGCTT |
| TEXEL1 deletion | 17 | PEXEL1_delete.F | CTCACAGAGAAAGTAAAAGTAGAGTTGCGTGA |
| | 18 | PEXEL1_delete.R | CCGCAGTCCGCCAATCGATC |
| Gra44 R1205A mutant generation (TEXEL1) | 19 | Quikchange R1205A.F | gattggcggactgcgggcagagctagaggaactc |
| | 20 | Quikchange R1205A.R | gagttcctctagctctgcccgcagtccgccaatc |
| Gra44 L1207 mutant generation (TEXEL1) | 21 | Quikchange L1207A.F | cggactgcggagagaggcagaggaactcacagag |
| | 22 | Quikchange L1207A.R | ctctgtgagttcctctgcctctctccgcagtccg |
| Gra44 E1209 mutant generation (TEXEL1) | 23 | Quikchange E1209A.F | gcggagagagctagaggcactcacagagaaagtaa |
| | 24 | Quikchange E1209A.R | ttactttctctgtgagtgcctctagctctctccgc |
| TEXEL2 deletion | 25 | PEXEL2_delete.F | TTGTTTGAACCGAAGAAAGAACCAAC |
| | 26 | PEXEL2_delete.R | CGAACCACTTCCTGCACG |
| Gra44 R1348A mutant generation (TEXEL2) | 27 | IMC2A-R1348A.F | AAGTGGTTCGgcCCGGCTCTTG |
| | 28 | IMC2A-R1348X.R | CCTGCACGTTTCTCCTCAGG |
| Gra44 L1350A mutant generation (TEXEL2) | 29 | IMC2A-L1350A.F | TTCGCGCCGGgcCTTGGAATT |
| | 30 | IMC2A-L1350X.R | CCACTTCCTGCACGTTTCTCCTC |
| Gra44 L1352A mutant generation (TEXEL2) | 31 | IMC2A E1352A.F | ccggctcttggcattgtttg |
| | 32 | IMC2A E1352X.R | cgcgaaccacttcctgc |
| Cloning TGGT1_316250 C-terminal region into endogenous tagging vector | 33 | 316250-myc.FOR | ttccaatccaatttaactgtgctaagccgttctagcgt |
| | 34 | 316250-myc.REV | ccacttccaattttatactgttccttagccatcatgtcgag |

Table S3. Primers used in this study.
